# Do clinical investigations predict long-term outcome? A follow-up of paediatric respiratory outpatients

**DOI:** 10.1101/529537

**Authors:** Carmen CM. de Jong, Eva SL. Pedersen, Myrofora Goutaki, Daniel Trachsel, Juerg Barben, Claudia E. Kuehni

## Abstract

**Introduction:** The contribution of clinical investigations to prediction of long-term outcomes of children investigated for asthma is unclear.

**Aim:** We performed a broad range of clinical tests and investigated whether they helped to predict long-term wheeze among children referred for evaluation of possible asthma.

**Methods:** We studied children aged 6-16 years referred to two Swiss pulmonary outpatient clinics with a history of wheeze, dyspnoea, or cough in 2007. The initial assessment included spirometry, body plethysmography, fractional exhaled nitric oxide, skin prick tests, and bronchial provocation tests (BPT) by exercise, methacholine, and mannitol. Respiratory symptoms were assessed with questionnaires at baseline and at follow-up seven years later. Associations between baseline factors and wheeze at follow-up were investigated by logistic regression.

**Results:** At baseline, 111 children were examined in 2007. Seven years after baseline, 85 (77%) completed the follow-up questionnaire, among whom 61 (72%) had wheeze at baseline, while at follow-up 39 (46%) reported wheeze. Adjusting for age and sex, the following characteristics predicted wheeze at adolescence: wheeze triggered by pets (odds ratio 4.2, 95% CI 1.2-14.8), pollen (2.8, 1.1-7.0), and exercise (3.1, 1.2-8.0). Of the clinical tests, only a positive exercise test (3.2, 1.1-9.7) predicted wheeze at adolescence.

**Conclusion:** Reported exercise-induced wheeze and wheeze triggered by pets or pollen were important predictors of wheeze persistence into adolescence. None of the clinical tests predicted wheeze more strongly than reported symptoms. Clinical tests might be important for asthma diagnosis but medical history is more helpful in predicting prognosis in children referred for asthma.

## Introduction

Asthma is the most prevalent chronic respiratory disease in childhood and adolescence, which leads to many health care visits.(1–3) Its key symptoms are wheeze, cough, and difficulty breathing, but symptoms vary substantially between individuals and across ages.(1,2) Some children who present with asthma symptoms continue to have problems later in life, while others do not. Better knowledge of their individual prognoses might affect their follow-up and answer questions of parents in the clinics.(4–6) Assessing prognosis of asthma symptoms from school age into adulthood and identifying children at high-risk of symptom persistence is challenging.(4)

Studies investigating prognosis of asthma or wheeze in school-aged children are conducted with either clinical asthma cohorts or symptomatic children of a population-based cohort.(7) Studies in clinical asthma cohorts have found that lower FEV1, asthma severity, sensitisation to indoor allergens, eczema, hay fever, skin test reactivity, and bronchial hyper-responsiveness were associated with asthma persistence.(8–10) Studies in population-based cohorts have found that wheeze persistence was predicted by frequent attacks of wheeze, female sex, sensitization to furred animals or house dust mites, rhinitis, and bronchial hyper-responsiveness.(11–16)

For clinical practice, two knowledge gaps remain. First, few studies have examined the prediction of long-term prognosis, but none have done this for school-aged children seen in outpatient clinics for possible asthma. Second, many tests are performed in clinics to diagnose these children, but it is unclear whether these tests predict prognosis more accurately than reported symptoms alone. We determined whether clinical tests in addition to reported symptoms help predict wheeze in adolescence in school-aged children referred for possible asthma.

## Methods

### Study population and study design

Of the 124 children invited, 111 were recruited from the respiratory outpatient clinics of two paediatric hospitals in Switzerland, 84 from St. Gallen and 27 from Basel, who were eligible if they had been referred for evaluation of current wheeze, dyspnoea, or cough. Children with a known chronic respiratory disease such as cystic fibrosis or primary ciliary dyskinesia, or a respiratory tract infection during four weeks prior to the visit were excluded. At baseline in 2007-2008, parents completed a questionnaire and children underwent a set of standardised clinical tests during two different visits within one week as part of the study protocol.(17, 18) At follow-up seven years after baseline, in 2014 to 2015, we sent a questionnaire to the 12-23 year-old adolescents or young adults (from now on referred to as adolescents) (Figure S2).

Ethical approval was obtained from the local Ethics committee and all parents gave informed consent at baseline (EKSG 07/001).

### Baseline assessment

The parental questionnaire included ISAAC key questions(19) plus additional questions on type and triggers of respiratory symptoms, atopic symptoms, previous treatments and environmental exposures. The study physician reported clinical test results, final diagnosis, and prescribed medication in a uniform way.

At the initial visit, children underwent performed spirometry, fractional exhaled nitric oxide (FeNO) measurement, a skin prick test (SPT), bronchial provocation test (BPT) by exercise and, by methacholine. At the second visit, children did a BPT by mannitol. Spirometry, and BPT by exercise and methacholine were performed according to published ATS guidelines.(20) A detailed description of the methodology of the clinical tests performed has been published elsewhere and is included in the online supplement. Lung function measurements were compared to reference values from Zapletal et al.(21) Details of the clinical tests are published elsewhere.(17) We considered the exercise test as positive in the event of a ≥15% decrease in the FEV1 after the exercise challenge test, and the methacholine test as positive when the minimal dose causing a ≥20% decrease of FEV1 was <1mg (the provocation dose, PD 20). The mannitol dry powder challenge test was considered as positive when a 15% fall in FEV1 was measured before a cumulative dose of 635 mg was reached, or when a 10% fall in FEV1 between two doses was reached. FeNO was measured using the portable NIOX MINO^®^ device, and was considered as positive when FeNO was higher than 26ppb.(18) We performed skin prick tests for birch, grass, mugwort, alternaria, cat, house dust mites (D. pteronysinus), and positive and negative controls.(18) These allergens cover 95% of inhaled allergens in Switzerland.(22) The test was considered to be positive if any mean wheal diameter was >3mm.

### Assessment at follow-up

The follow-up questionnaire was very similar to the baseline questionnaire, but the questions were addressed directly to the adolescents instead of their parents.

### Definitions of wheeze and frequent wheeze

We assessed wheeze at follow-up with the question, “Have you had a whistling sound in the chest in the last 12 months?” If a child had had more than three attacks of wheeze in the last 12 months, we considered the child to have had frequent wheeze.

### Statistical analysis

We compared the participants with information at baseline and follow-up to those without follow-up information to test for selection bias, using chi-square test. The participants with information at baseline and follow-up were included in the analysis.

We investigated the association between exercise-induced wheeze and a positive exercise test at baseline using the Fisher’s exact test, and the Mann-Whitney-U test when looking at the association of reported exercise-induced wheeze and the fall of FEV1% predicted during the exercise test.

We investigated the association between symptoms and clinical test results at baseline with any wheeze and frequent wheeze at follow-up using logistic regression. We included all symptoms from table 1 and clinical tests from table 2 in the model and adjusted for sex and age. We did not consider interactions because of the sample size. We used STATA software (version 14; College Station, Texas) to analyse the data.

## Results

### Characteristics of the study population at baseline and at follow-up

Eighty-five (77%) of the 111 children who participated in the baseline study completed the follow-up questionnaire. The median age was 12 years at baseline (range 6-16) and 18 at follow-up (12-23); 60% (51/85) were male. Wheeze was reported by 61 (72%) at baseline, and 7 reported cough without wheeze and 12 (14%) reported exercise-related breathing problems. Among those with wheeze, 27 (44%) had more than three attacks during 12 months prior to the baseline visit (Table 1). Symptoms at baseline were very similar in children who did not take part in the follow-up (Table S1 online supplementary material). Asthma medication was prescribed at the baseline visit for 71 (85%) children, of whom 47 (55%) received inhaled short-acting β2-agonists (SABA) alone, 6 received SABA and inhaled corticosteroids (ICS), and 18 received long-acting β2-agonists (LABA) and ICS. At follow-up, 39 (46%) participants reported wheeze of whom 30 had more than 3 attacks during the last year. At follow-up, 44 adolescents (52%) reported using inhalers, including 21 using SABA alone, 2 using SABA and ICS, and 21 using LABA and ICS (Table 1).

**Table 1:**
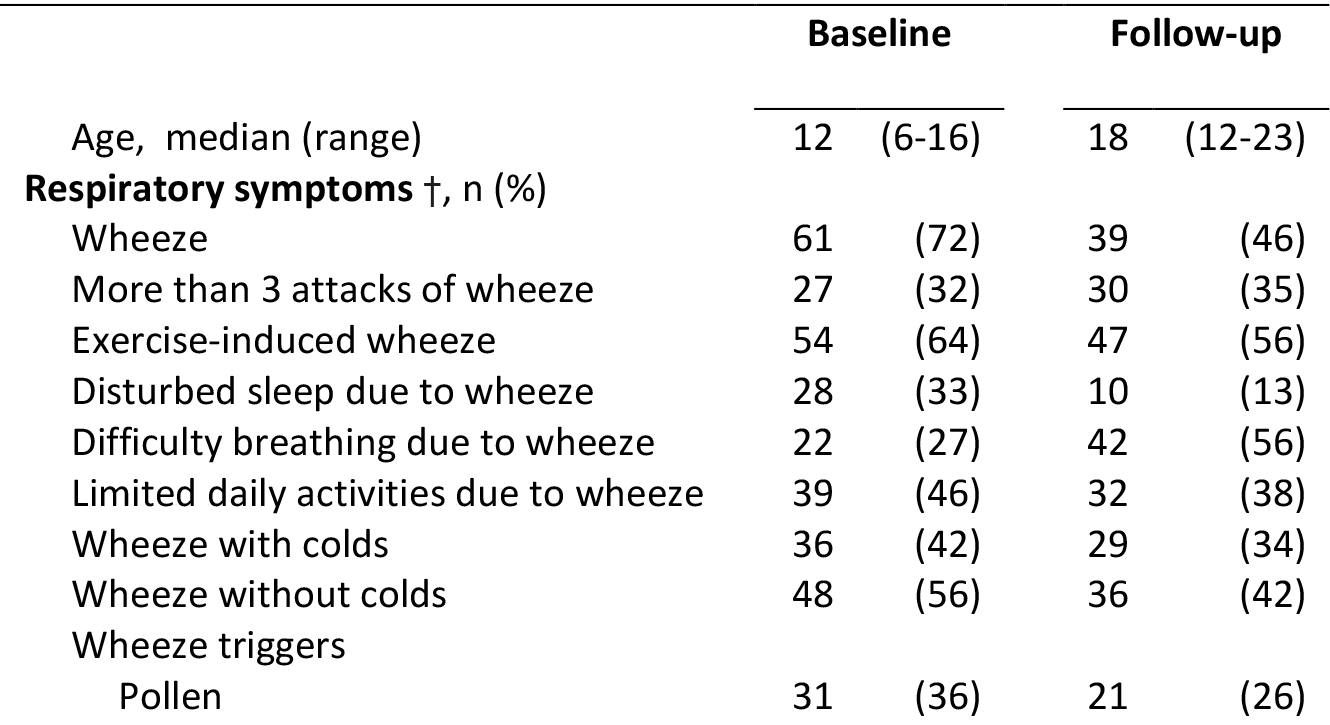
Characteristics of the study population at baseline and follow-up (N=85)

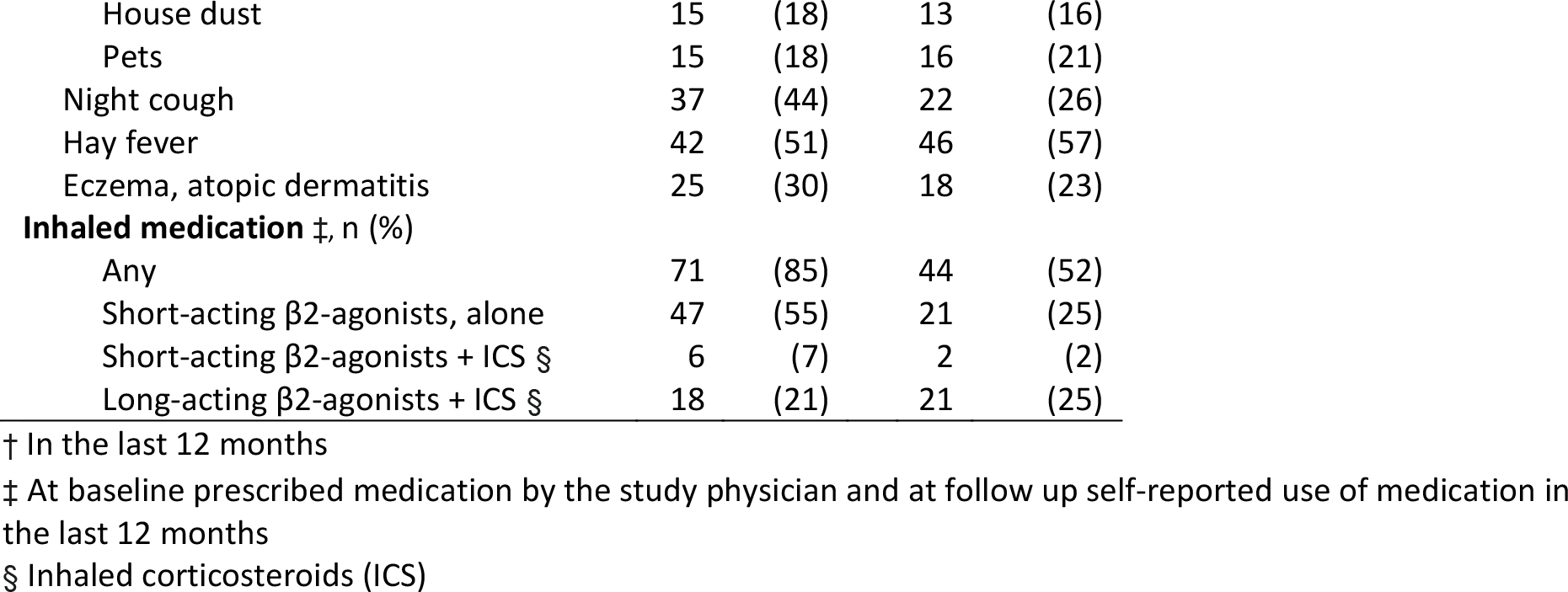

Table 2 shows the clinical test results and diagnoses at baseline. All tests were completed in at least 90% of the children. The main reason for not completing a BPT was exhaustion.(17,18) For the 78 children who completed the BPT by methacholine at baseline, the test was positive in 76% and the median provocation dose was 0.14mg. Eighty-two completed the BPT by mannitol, of whom 28% tested positive. The median provocation dose was 635 mg. Of the 76 children who completed the BPT by exercise, the test was positive in 24% with a median fall of FEV1 of 8% predicted. SPT was positive in 33 (39%) children and FeNO was positive in 35 (41%). Doctors diagnosed 62 (73%) children with asthma or episodic viral wheeze. The other children were mostly diagnosed with cough not due to asthma or vocal cord dysfunction.

At baseline, self-reported exercise-induced wheeze was associated with a positive exercise test (p=0.022, table S2). Self-reported exercise-induced wheeze was also associated with the fall of FEV1% predicted during the exercise test (p=0.003, figure S1)

**Table 2.**
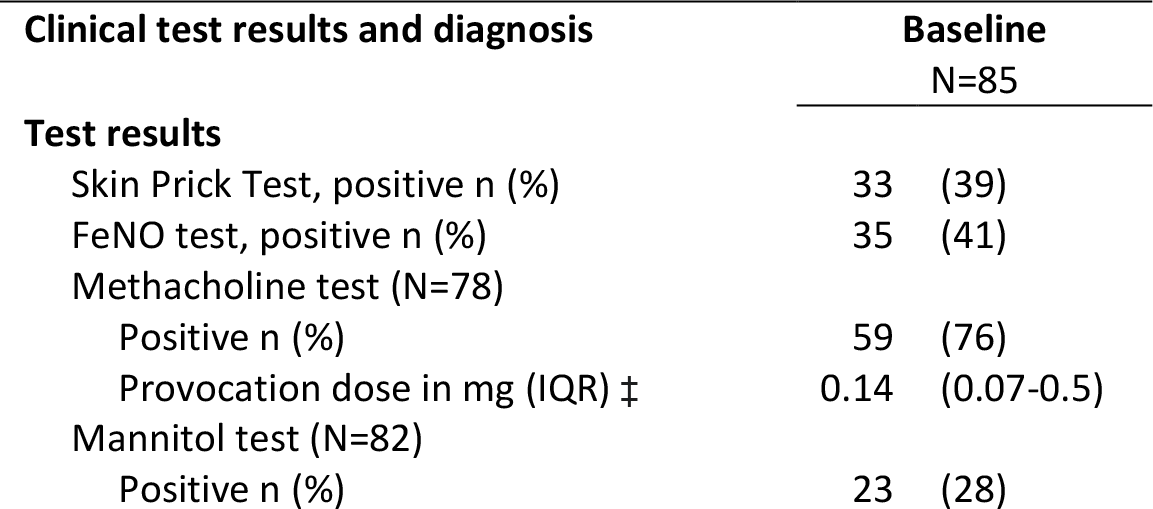
Results of clinical tests and final diagnosis at baseline

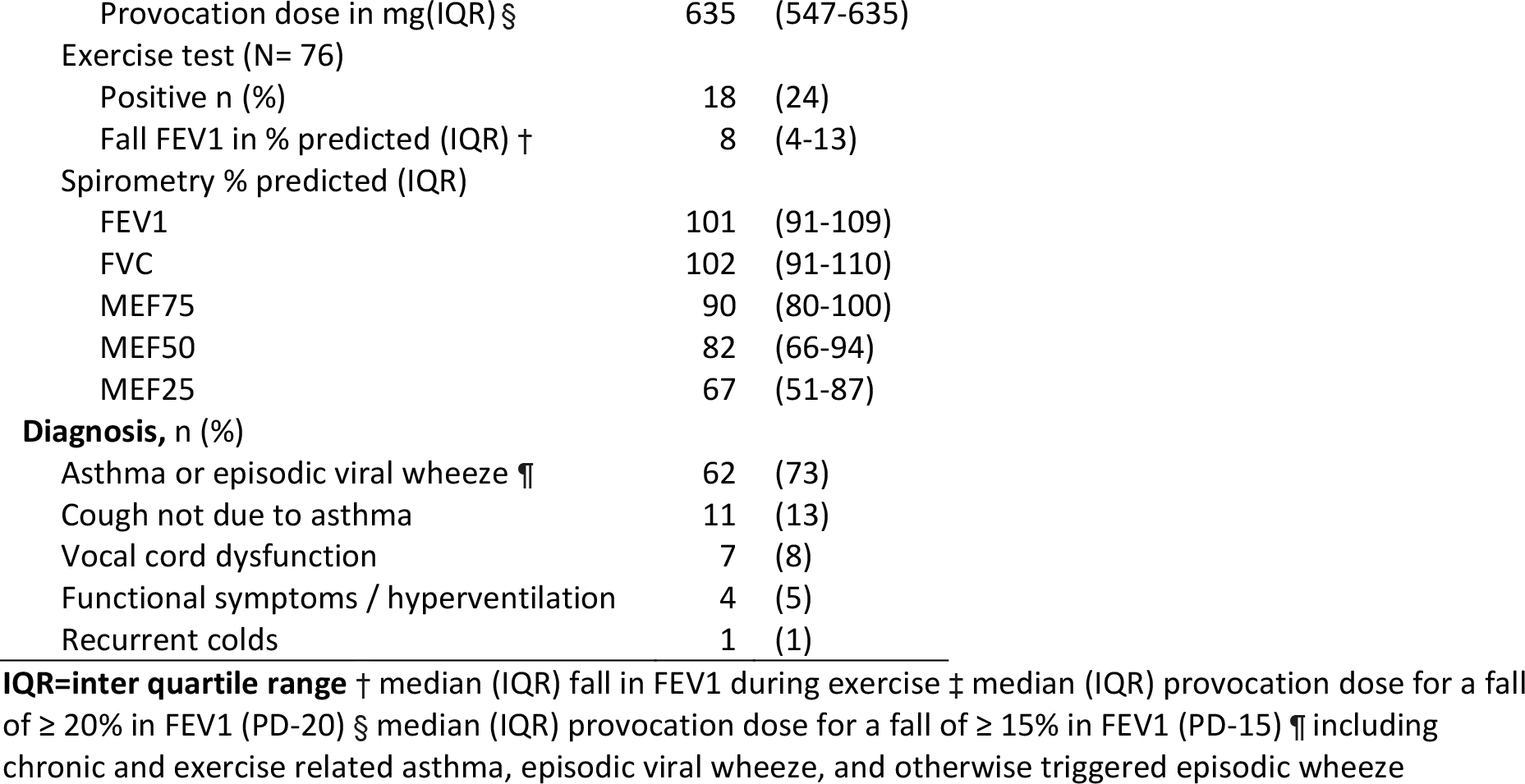

### Baseline factors associated with wheeze and frequent wheeze at follow-up

Four respiratory symptoms and one clinical test at baseline were associated with *any wheeze* at follow-up. Of the reported symptoms, frequent wheeze (>3 attacks) (OR 2.86, 95% CI 1.10-7.43), exercise-induced wheeze (3.07, 1.19-7.96), wheeze triggered by pets (4.22, 1.21-14.76), and wheeze triggered by pollen (2.78, 1.11-6.98) were associated with wheeze at follow-up. For the clinical tests, only a positive exercise test was significantly associated with wheeze seven years later (3.20, 1.05-9.70). Results remained very similar after adjusting for age and sex (Table 3, Fig. 1).

Two respiratory symptoms were associated with *frequent wheeze* at follow-up. These were exercise-induced wheeze (OR 3.05, 95% CI 1.07-8.67) and wheeze triggered by pets (3.79, 1.15-12.48; Table 4). None of the clinical test results were associated with frequent wheeze at follow-up.

**Table 3:**
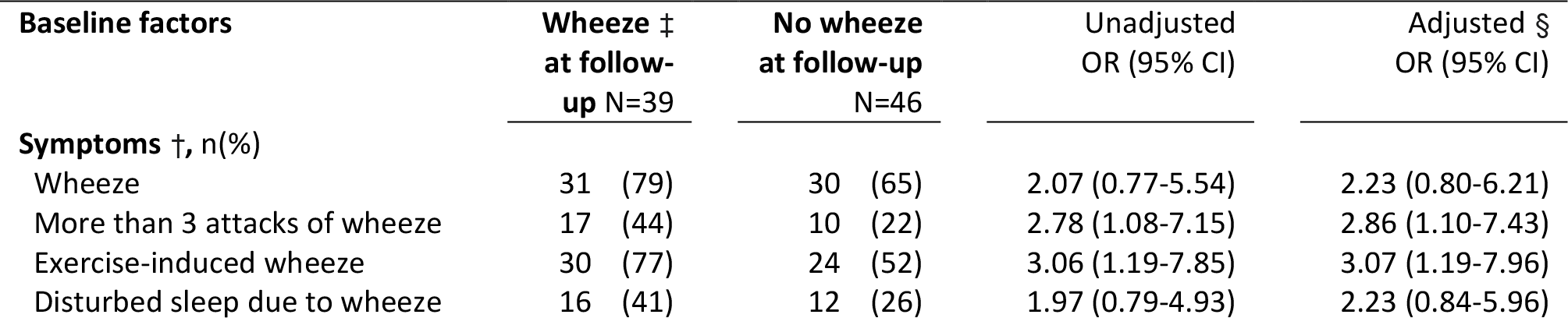
Associations between baseline factors and wheeze at follow up

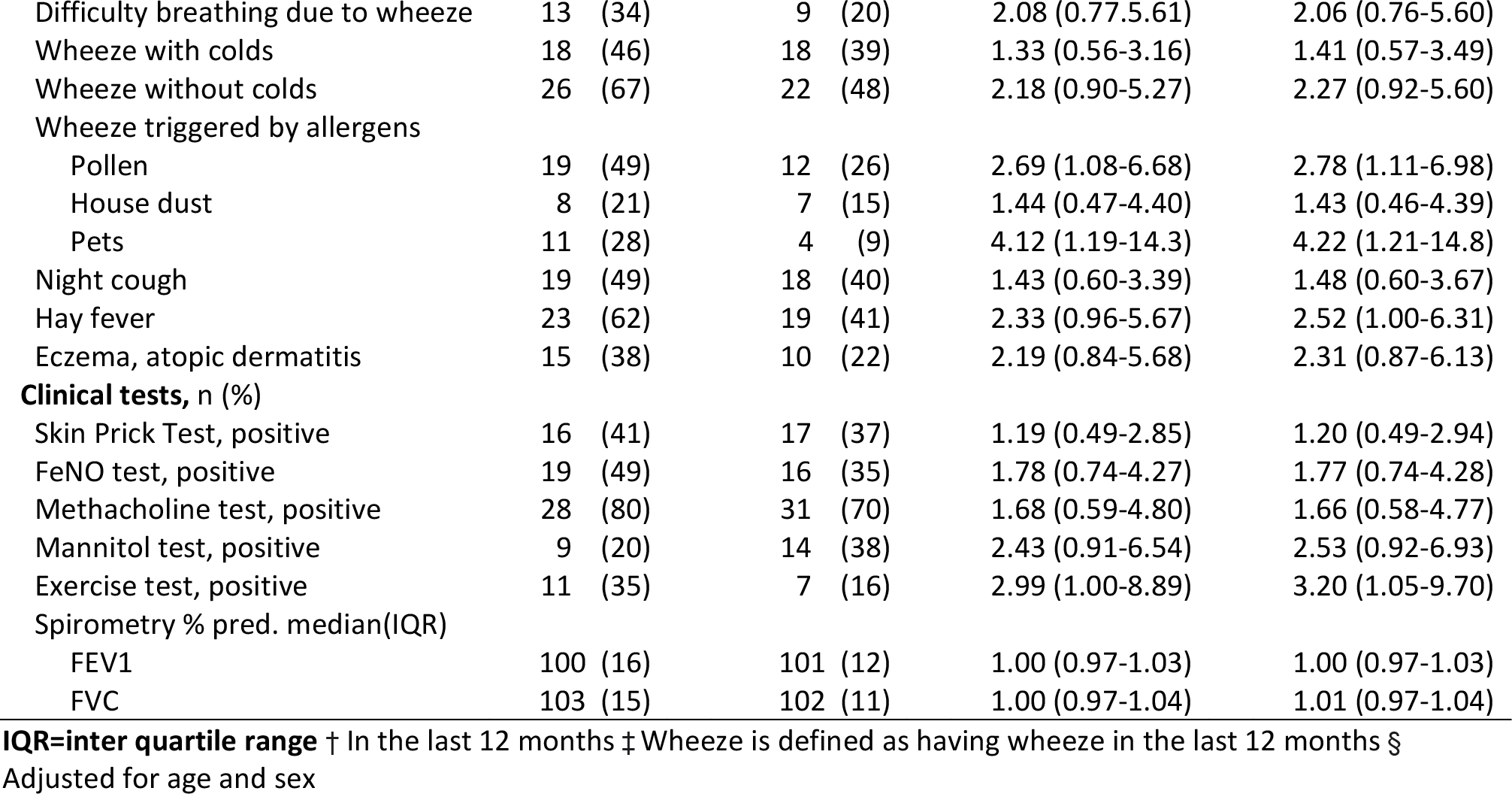

**Figure 1:**
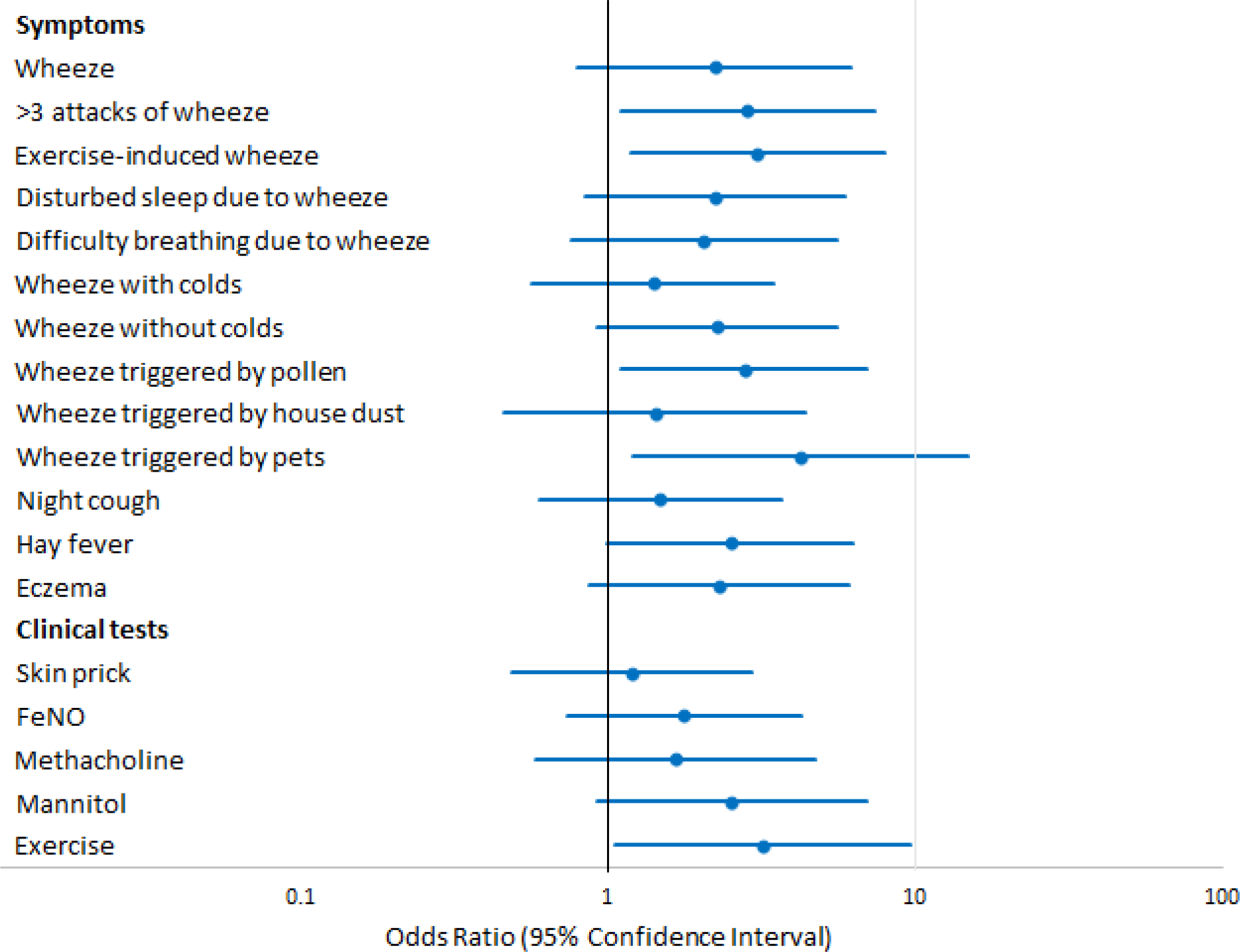
Associations between baseline factors and wheeze at follow up.

**Table 4:**
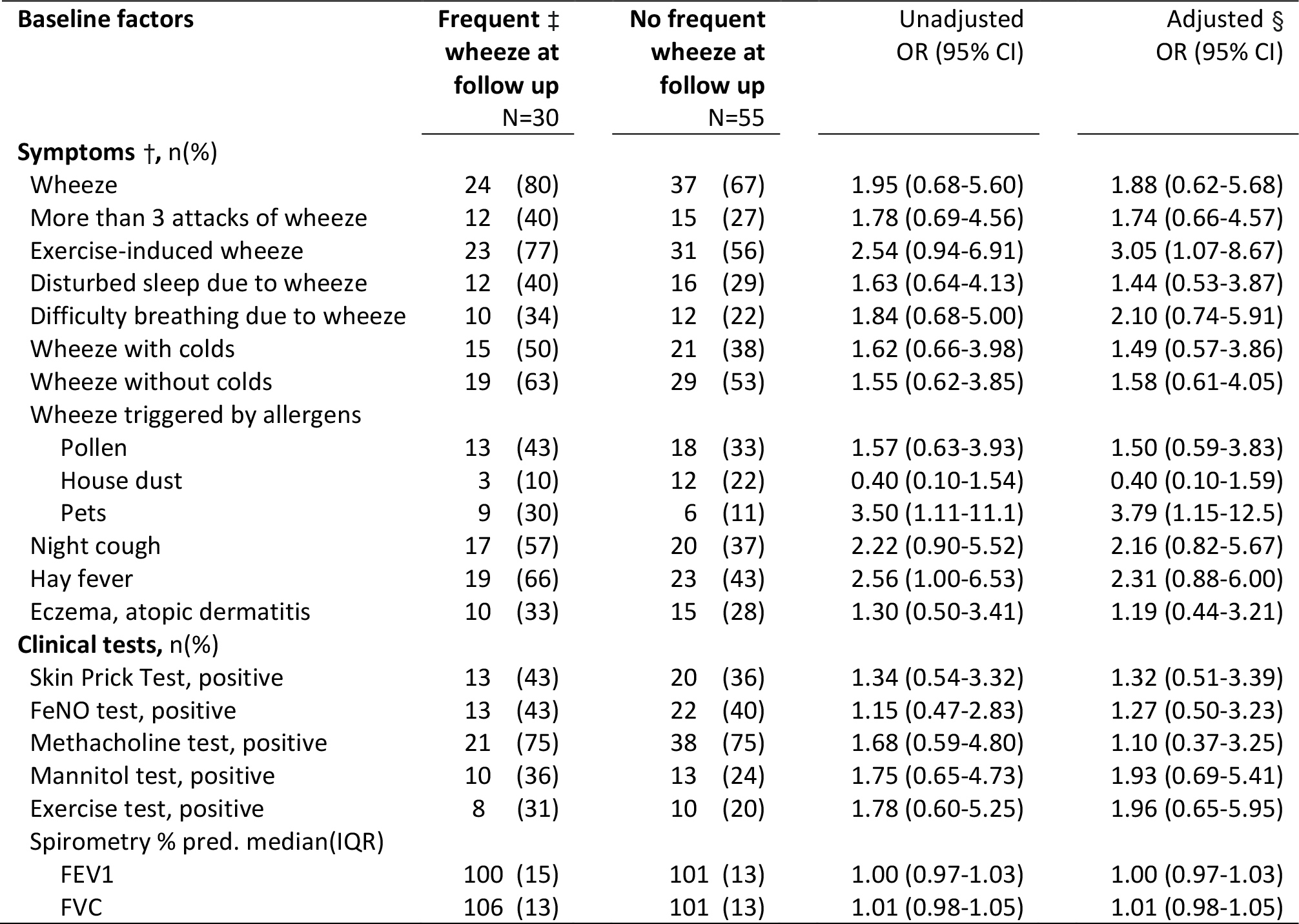
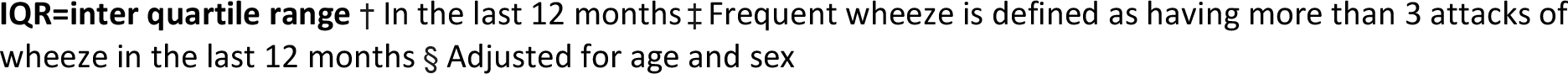
Associations between baseline factors and frequent wheeze at follow up

| Baseline factors | Frequent ‡<br>wheeze at<br>follow up<br>N=30 | No frequent<br>wheeze at<br>follow up<br>N=55 | Unadjusted<br>OR (95% CI) | Adjusted §<br>OR (95% CI) |
| --- | --- | --- | --- | --- |
| **Symptoms †, n(%)** |  |  |  |  |
| Wheeze | 24 (80) | 37 (67) | 1.95 (0.68-5.60) | 1.88 (0.62-5.68) |
| More than 3 attacks of wheeze | 12 (40) | 15 (27) | 1.78 (0.69-4.56) | 1.74 (0.66-4.57) |
| Exercise-induced wheeze | 23 (77) | 31 (56) | 2.54 (0.94-6.91) | 3.05 (1.07-8.67) |
| Disturbed sleep due to wheeze | 12 (40) | 16 (29) | 1.63 (0.64-4.13) | 1.44 (0.53-3.87) |
| Difficulty breathing due to wheeze | 10 (34) | 12 (22) | 1.84 (0.68-5.00) | 2.10 (0.74-5.91) |
| Wheeze with colds | 15 (50) | 21 (38) | 1.62 (0.66-3.98) | 1.49 (0.57-3.86) |
| Wheeze without colds | 19 (63) | 29 (53) | 1.55 (0.62-3.85) | 1.58 (0.61-4.05) |
| Wheeze triggered by allergens |  |  |  |  |
| Pollen | 13 (43) | 18 (33) | 1.57 (0.63-3.93) | 1.50 (0.59-3.83) |
| House dust | 3 (10) | 12 (22) | 0.40 (0.10-1.54) | 0.40 (0.10-1.59) |
| Pets | 9 (30) | 6 (11) | 3.50 (1.11-11.1) | 3.79 (1.15-12.5) |
| Night cough | 17 (57) | 20 (37) | 2.22 (0.90-5.52) | 2.16 (0.82-5.67) |
| Hay fever | 19 (66) | 23 (43) | 2.56 (1.00-6.53) | 2.31 (0.88-6.00) |
| Eczema, atopic dermatitis | 10 (33) | 15 (28) | 1.30 (0.50-3.41) | 1.19 (0.44-3.21) |
| **Clinical tests, n(%)** |  |  |  |  |
| Skin Prick Test, positive | 13 (43) | 20 (36) | 1.34 (0.54-3.32) | 1.32 (0.51-3.39) |
| FeNO test, positive | 13 (43) | 22 (40) | 1.15 (0.47-2.83) | 1.27 (0.50-3.23) |
| Methacholine test, positive | 21 (75) | 38 (75) | 1.68 (0.59-4.80) | 1.10 (0.37-3.25) |
| Mannitol test, positive | 10 (36) | 13 (24) | 1.75 (0.65-4.73) | 1.93 (0.69-5.41) |
| Exercise test, positive | 8 (31) | 10 (20) | 1.78 (0.60-5.25) | 1.96 (0.65-5.95) |
| Spirometry % pred. median(IQR) |  |  |  |  |
| FEV1 | 100 (15) | 101 (13) | 1.00 (0.97-1.03) | 1.00 (0.97-1.03) |
| FVC | 106 (13) | 101 (13) | 1.01 (0.98-1.05) | 1.01 (0.98-1.05) |

**IQR=inter quartile range** † In the last 12 months‡ Frequent wheeze is defined as having more than 3 attacks of wheeze in the last 12 months§ Adjusted for age and sex

## Discussion

Among school-aged children referred to a respiratory outpatient clinic for evaluation of wheeze, cough, or dyspnoea, 46% reported wheeze seven years later. Reported exercise-induced wheeze and wheeze triggered by pets or pollen at baseline predicted wheeze at follow-up. Of the clinical tests, only a positive exercise challenge test predicted wheeze at follow-up, but no more strongly than reported exercise-induced wheeze.

A few studies have examined the prediction of prognosis by clinical testing, but ours is the only study to have done this for so many clinical tests in school-aged children referred to a respiratory outpatient clinic. We did not find an association between FEV1 at baseline and wheeze seven years later; previous studies have reported contradictory findings. Both the CAMP cohort of 909 children aged 5-12 years with diagnosed asthma and another Dutch clinical cohort study of 5-14 year-old children diagnosed with asthma found that asthma persistence at ages 15-20 and 32-42, respectively, was associated with decreased FEV1 at school-age.(8, 9) In contrast, the population-based Tasmanian cohort did not find an association between FEV1 at age 7 and wheeze persistence at age 29-32.(12, 15) This could be because children with wheeze from population-based cohorts might have milder asthma than those in clinical studies. Our findings are not directly comparable with those of other studies that studied younger children, reported test results differently, or included healthy children.

Our observation that frequent attacks of wheeze at school age predicted wheeze persistence seven years later is in line with findings from the Melbourne and Tasmanian cohorts.(10, 11) In contrast to their findings, we found no significant association between either eczema or hay fever at baseline and wheeze persistence. This could be because those cohorts used different outcomes—severe wheeze and atopic asthma, respectively—or simply because we had low numbers and limited power.

A possible limitation of our study was that the bronchial provocation tests were done within a short period of time. This could have influenced the methacholine test result, which was performed after the exercise test on the same day and was positive in 76% of the children. Most likely the bronchial provocation test by mannitol was not influenced by the short time interval. We assured an appropriate interval of at least 24 hours without a change in respiratory health or medication in this time interval. A second limitation was the small sample size, which limited statistical power and did not allow us to perform a multivariable analysis including all symptoms and test results simultaneously.

The main strength of our study is its clinical design, which reflects the typical mix of patients in a paediatric outpatient clinic. All children were first-time referrals to the paediatric respiratory clinic for evaluation of possible asthma. Therefore, the study population is representative of daily clinical work, in contrast to many clinical studies that selectively include well-defined moderate to severe asthmatics and leave out patients with unclear degrees of airway reactivity. Our study also profited from a very detailed baseline examination. Children in the study had an extensive array of examinations for lung function, BPT and allergy, which allowed us to assess the contribution of clinical tests in predicting long-term wheeze in addition to reported symptoms among those referred for evaluation of possible asthma.

## Conclusion

This study is an initial step towards finding out whether clinical tests can predict wheeze later in life. Though clinical tests might be important for asthma diagnosis, our results suggest that they do not strongly predict prognosis of wheeze. In contrast, our data underline the importance of a detailed history, as school-age children reporting exercise-related wheeze and wheeze triggered by allergens were at higher risk and thus might profit from more frequent follow-up.

## General acknowledgements

We thank all participants and lab technicians of the pulmonology department in the children’s hospitals in Basel and St. Gallen for their assistance in our study, Marie-Pierre Strippoli (ISPM, Bern) for her work on the study at baseline, Bettina Meier (ISPM, Bern) for entering the follow-up questionnaires into the database and sending the participation reminders. We thank Christopher Ritter (ISPM, Bern) for his editorial assistance and Niels Hagenbuch (ISPM, Bern) for his statistical support.

## Author contributions

Claudia Kuehni and Jürg Barben conceptualised and designed the study. Daniel Trachsel and Jürg Barben supervised data collection. Carmen de Jong analysed the data and drafted the manuscript. Eva Pedersen and Myrona Goutaki supported the statistical analysis and gave input for interpretation of the data. All authors critically revised the manuscript and approved the final manuscript as submitted.

## Supplementary data

**Table S1.** Comparison of characteristics of the children included in the follow-up study and the children that did not take part in the follow-up study

**Table S2** Association between reported exercise-induced wheeze and exercise test result at baseline N=76

**Figure S1:** Association between reported exercise-induced wheeze and the fall of FEV1% predicted during exercise testing at baseline

