## Supplementary Table 1 and 2 for "Do clinical investigations predict long-term outcome? A follow-up of paediatric respiratory outpatients"

**Table S1.** Comparison of characteristics of the children included in the follow-up study and the children that did not take part in the follow-up study

| Characteristics | Complete information, Included N=85 | | No follow-up information, Not included N=27 | | p-value″ |
|---|---|---|---|---|---|
| Age, median (range) | 12 | (6-16) | 11 | (6-15) | 0.269 |
| Sex, male n(%) | 8 | (31) | 34 | (40) | 0.396 |
| **Respiratory symptoms***, n(%) | | | | | |
| Wheeze | 61 | (72) | 19 | (73) | 0.896 |
| More than 3 attacks of wheeze | 27 | (32) | 11 | (42) | 0.321 |
| Exercise-induced wheeze | 54 | (64) | 16 | (62) | 0.854 |
| Disturbed sleep due to wheeze | 28 | (33) | 8 | (31) | 0.836 |
| Difficulty breathing due to wheeze | 22 | (27) | 3 | (12) | 0.113 |
| Limited daily activities due to wheeze | 39 | (46) | 12 | (46) | 0.981 |
| Wheeze with colds | 36 | (42) | 7 | (26) | 0.126 |
| Wheeze without colds | 48 | (56) | 19 | (70) | 0.199 |
| Wheeze triggers | | | | | |
| Exercise | 54 | (64) | 15 | (56) | 0.458 |
| Laughing | 11 | (13) | 5 | (19) | 0.471 |
| Pollen | 31 | (36) | 5 | (19) | 0.082 |
| House dust | 15 | (18) | 6 | (22) | 0.596 |
| Pets | 15 | (18) | 5 | (26) | 0.697 |
| Food/drinks | 3 | (4) | 0 | (0) | 0.322 |
| Night cough | 37 | (44) | 11 | (42) | 0.876 |
| Hay fever | 42 | (51) | 7 | (27) | 0.034 |
| Eczema, atopic dermatitis | 25 | (30) | 1 | (4) | 0.008 |
| Parental smoking | 23 | (27) | 8 | (30) | 0.795 |
| **Inhaled Medication**#, n(%) | | | | | |
| Any | 71 | (85) | 15 | (58) | 0.004 |
| Short-acting β2-agonists | 47 | (55) | 7 | (26) | 0.008 |
| Short-acting β2-agonists + ICS° | 6 | (7) | 3 | (11) | 0.500 |
| Long-acting β2-agonists + ICS° | 18 | (21) | 5 | (19) | 0.766 |

* In the last 12 months # At baseline prescribed medication by the study physician after the diagnostic tests and at follow up self-reported use of medication in the last 12 months ° Inhaled corticosteroids (ICS) “ chi-square test for dichotomous and t-test for continuous variables.

**Table S2** Association between reported exercise-induced wheeze and exercise test result at baseline N=76

| Reported exercise-induced wheeze | Exercise test negative n(%column)[%row] | Exercise test positive n(%column)[%row] | Total |
|---|---|---|---|
| No | 25 (43) [93] | 2 (11) [7] | 27 |
| Yes | 33 (57) [67] | 16 (89) [33] | 49 |
| **Total** | 58 | 18 | |

Fisher's exact: p-value 0.022
