## Supplementary Figure 1 for "Do clinical investigations predict long-term outcome? A follow-up of paediatric respiratory outpatients"

**Figure S1:** Association between reported exercise-induced wheeze and the fall of FEV1% predicted during exercise testing at baseline

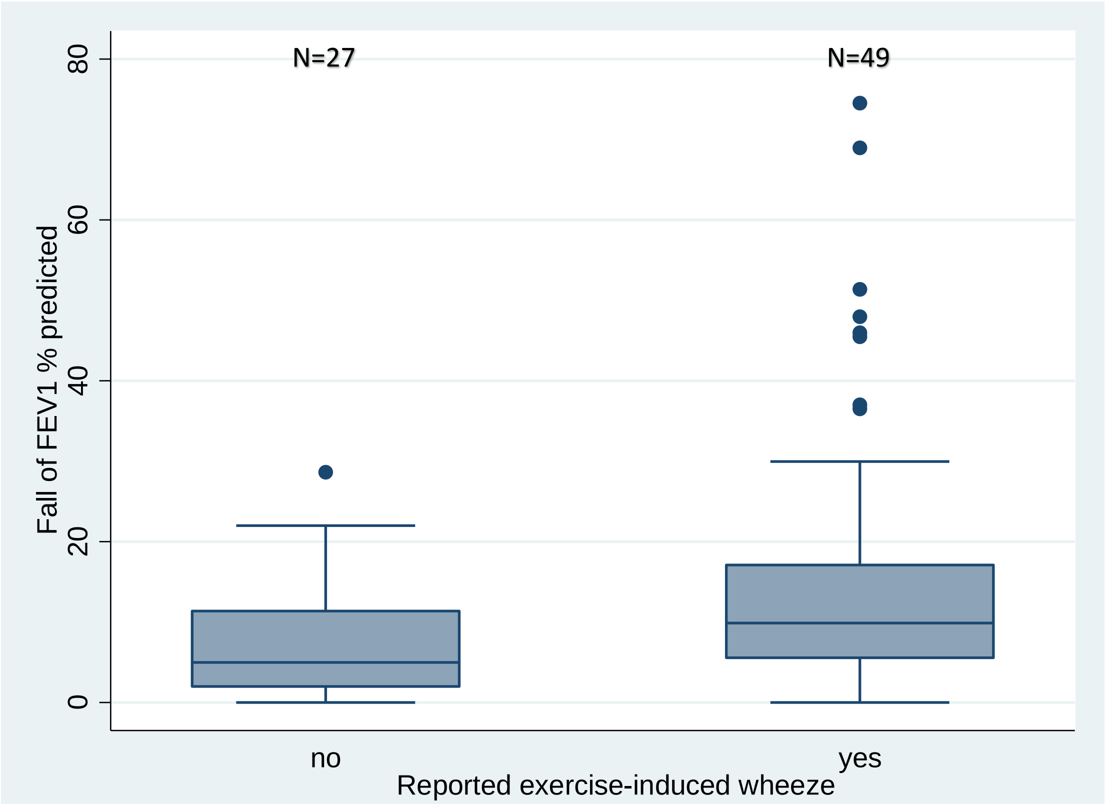


Mann-Whitney-U: p-value 0.003
